# A Neutralizing Antibody-Conjugated Photothermal Nanoparticle Captures and Inactivates SARS-CoV-2

**DOI:** 10.1101/2020.11.30.404624

**Authors:** Xiaolei Cai, Aleksander Prominski, Yiliang Lin, Nicholas Ankenbruck, Jillian Rosenberg, Min Chen, Jiuyun Shi, Eugene B. Chang, Pablo Penaloza-MacMaster, Bozhi Tian, Jun Huang

## Abstract

The outbreak of 2019 coronavirus disease (COVID-19), caused by severe acute respiratory syndrome coronavirus 2 (SARS-CoV-2), has resulted in a global pandemic. Despite intensive research including several clinical trials, currently there are no completely safe or effective therapeutics to cure the disease. Here we report a strategy incorporating neutralizing antibodies conjugated on the surface of a photothermal nanoparticle to actively capture and inactivate SARS-CoV-2. The photothermal nanoparticle is comprised of a semiconducting polymer core and a biocompatible polyethylene glycol surface decorated with neutralizing antibodies. Such nanoparticles displayed efficient capture of SARS-CoV-2 pseudoviruses, excellent photothermal effect, and complete inhibition of viral entry into ACE2-expressing host cells via simultaneous blocking and inactivating of the virus. This photothermal nanoparticle is a flexible platform that can be readily adapted to other SARS-CoV-2 antibodies and extended to novel therapeutic proteins, thus providing a broad range of protection against multiple strains of SARS-CoV-2.

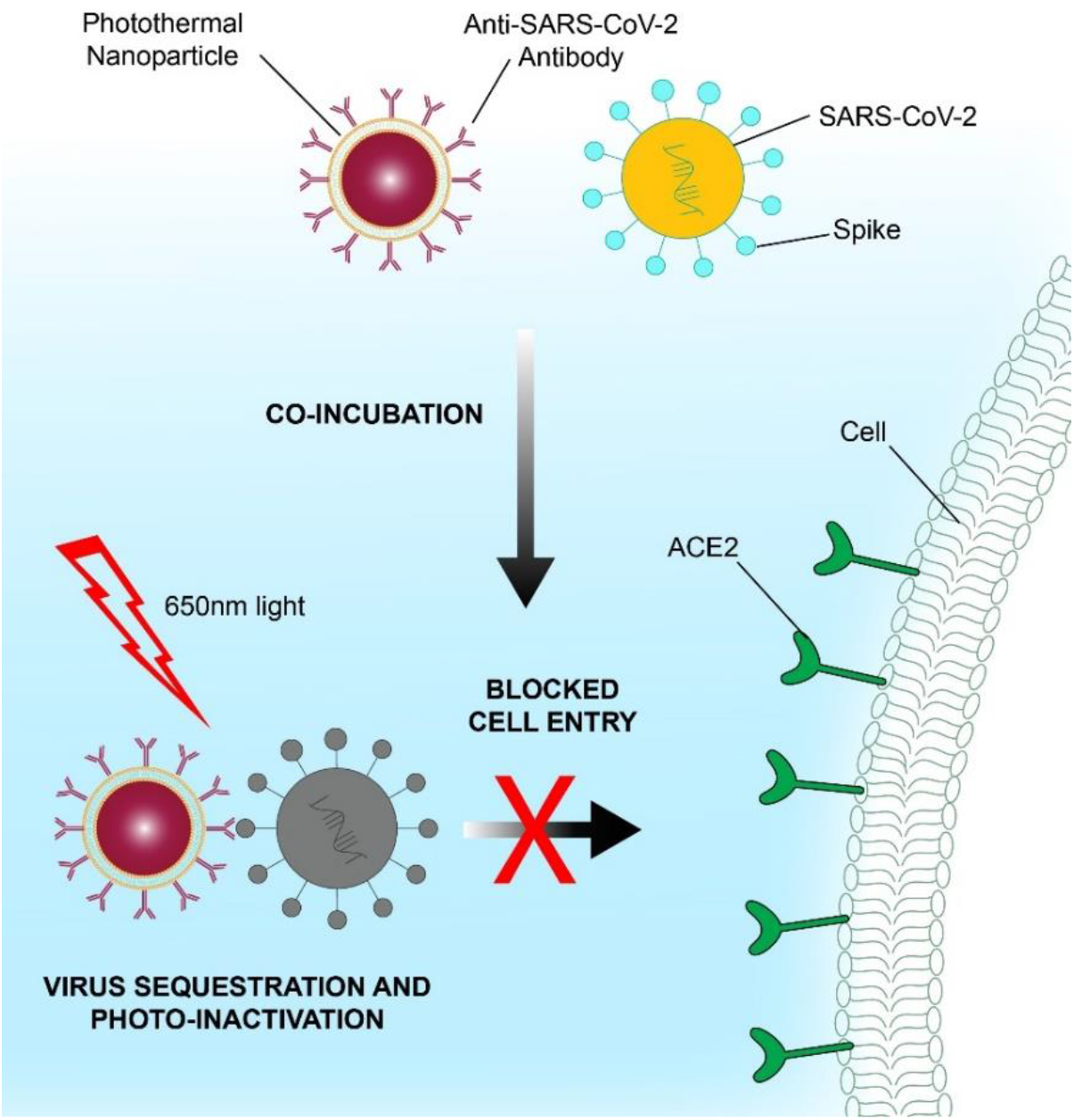

---

Coronavirus disease 2019 (COVID-19), resulting from severe acute respiratory syndrome coronavirus 2 (SARS-CoV-2) infection, has spread worldwide and caused a global pandemic.^1, 2^ As of October 12, 2020, SARS-CoV-2 has infected over 37 million people and caused more than one million deaths.^3^ The virus can induce severe symptoms such as acute respiratory distress syndrome, cytokine release syndrome, and cardiovascular damage, as well as increase mortality in patients.^4–6^ SARS-CoV-2 gains entry into cells through high affinity binding of the receptor binding domain of the spike protein to the angiotensin-converting enzyme 2 (ACE2) receptor on the host cell surface.^7, 8^

Vaccines in development to protect against SAR-CoV-2 infection have shown promising results in phase 1 and 2 clinical trials,^9–13^ however the process to vaccine approval and deployment may take months to years to realize. Moreover, it remains unclear whether vaccines will confer longterm protection, further complicating the path toward ending the pandemic.^14^ Currently, there are limited therapeutic regimens proven to clear the viral infection in all patients evaluated. Existing FDA-approved drugs have been applied to COVID-19 treatment and shown some success, but specific, effective therapies have been yet to be clinically approved.^15–17^ In an alternative approach, transfusion of convalescent plasma from recovered patients containing antibodies specific for SARS-CoV-2 into COVID-19 patients was shown to reduce viral load and may limit the severity or duration of illness in some patients due to the presence of pre-existing neutralizing antibodies.^18^ However, intravenous administration of convalescent plasma involves logistical hurdles, including availability of donor plasma and the need for a designated medical facility.^19^

Antibody-dependent enhancement (ADE), caused by binding of non-neutralizing antibodies to SARS-CoV-2, remains a concern for antibody-based vaccines, therapies, and convalescent plasma because it can further aggravate a patient’s condition through enhancement of viral infection.^20–23^ High-affinity monoclonal neutralizing antibodies against SARS-CoV-2 may circumvent some of the potential risk of ADE, as they often display much higher affinities for SARS-CoV-2 spike protein (<1 nM)^24–26^ than those of ACE2 (~15–40 nM)^27–30^ and can be produced in mass scale. Recently, several monoclonal neutralizing antibodies cloned using B cells from COVID-19 patients have been shown to effectively block the interaction between SARS-CoV-2 spike protein and ACE2.^31–35^

Several nanomaterial-based approaches have been investigated for virus detection,^36–39^ vaccine delivery,^40, 41^ and viral capture,^42–44^ however, the lack of a method to effectively capture and inactivate the virus after binding—which may lead to ADE—remains to be addressed. Herein we report the development of a strategy utilizing photothermal nanoparticles decorated with high-affinity neutralizing antibodies in order to effectively capture and inactivate SARS-CoV-2 (Figure 1a). Each photothermal nanoparticle contains a semiconducting polymer core, poly[2,6-(4,4-bis-(2-ethylhexyl)-4H-cyclopenta [2,1-b;3,4-b’]dithiophene)-alt-4,7(2,1,3-benzothiadiazole)] (PCPDTBT), enabling generation of intensive local heat after being excited by suitable light sources. In addition, photothermal nanoparticles can achieve highly specific and noninvasive photothermal killing of biological targets such as virus, bacteria, and tumors without damaging the surrounding healthy tissues.^45–47^ An amphiphilic polymer shell, 2-distearoyl-sn-glycero-3-phosphoethanolamine-N-[methoxy(polyethylene glycol)-2000], is applied to encapsulate the PCPDTBT core. The nanoparticle surface is functionalized with a monoclonal neutralizing antibody specific to the SARS-CoV-2 spike protein which enables selective and efficient capturing of SARS-CoV-2 with high affinity (0.07 nM), thus preventing the entry of SARS-CoV-2 into host cells. Upon excitation by a 650-nm light-emitting diode (LED), which possesses a more desirable safety profile compared to conventional laser excitation,^48, 49^ the photothermal nanoparticles directly inactivate the captured SARS-CoV-2 by heat. The unique design of our photothermal nanoparticles not only synergizes with the neutralizing function of the antibody in capturing SARS-CoV-2, but may also mitigate the potential risks of ADE through direct inactivation of the coronavirus via the photothermal effect.

**Figure 1.**
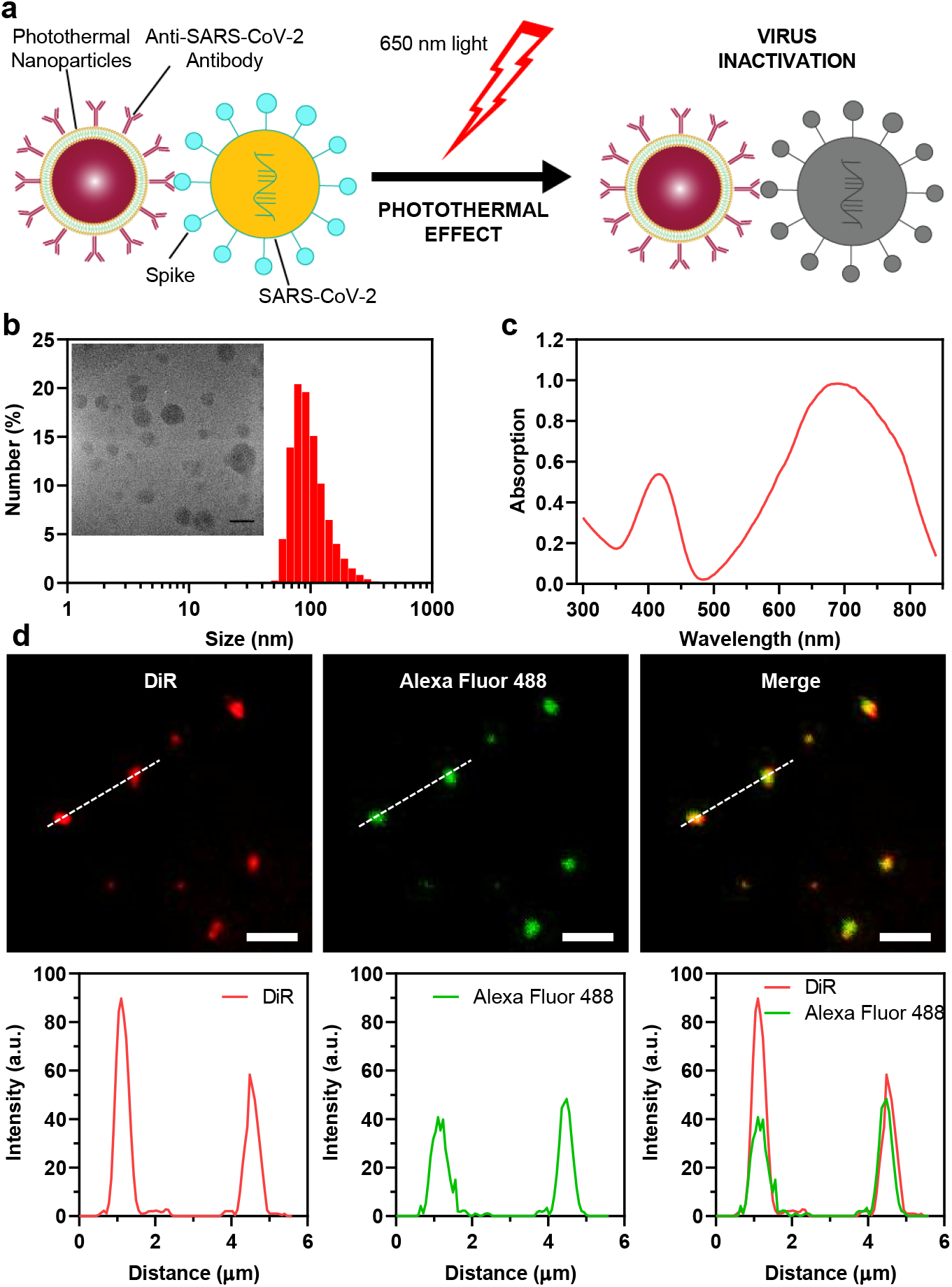
Structure and characterization of photothermal nanoparticles. **(a)** Schematic illustration of the photothermal nanoparticle for virus capture and inactivation. **(b)** DLS size distribution of photothermal nanoparticles. Inset: a TEM image of the photothermal nanoparticles. Scale bar: 50 nm. **(c)** UV-vis absorption spectrum of photothermal nanoparticles. **(d)** Fluorescent images and fluorescence intensity profiles of the neutralizing antibody-conjugated photothermal nanoparticles labeled by DiR (red) and immunostaining by a secondary Alexa Fluor-488 anti-IgG2b antibody (green) and their overlaid images. Line scan was used to indicate the fluorescence profile and co-localization for single nanoparticles. Excitation: 488 nm (green channel) and 740 nm (red channel). Scale bars: 2.5 μm.

## RESULTS AND DISCUSSION

Photothermal nanoparticles were prepared by using 1,2-distearoyl-sn-glycero-3-phosphoethanolamine-N-[carboxy(polyethylene glycol)-2000, NHS ester] (DSPE-PEG2000-NHS) as the matrix to encapsulate PCPDTBT through self-assembly.^47^ Subsequently, an anti-SARS-CoV-2 neutralizing antibody was crosslinked to the nanoparticles via reaction of NHS esters with amines on the antibody surface. Using dynamic light scattering (DLS), we determined the hydrodynamic diameters of the photothermal nanoparticles to be ~ 90 nm, which was validated by transmission electron microscopy (TEM) (Figure 1b). In addition, the photothermal nanoparticles exhibited good stability without forming any aggregation or precipitation after being stored in aqueous dispersions at 4 °C for several weeks (Figure S1). The ultraviolet to the visible light absorption spectrum of the photothermal nanoparticles covers a broad range from 500 to 850 nm in the red and near-infrared range, favoring their application as photothermal agents for light-triggered inactivating of the SARS-CoV-2 using infrared excitation (Figure 1c). Surface conjugation of the anti-SARS-CoV-2 neutralizing antibody (IgG2b) was further validated by fluorescent imaging analysis. Briefly, DiOC18(7) (DiR) lipophilic dye capable of emitting a fluorescent signal at ~780 nm was co-encapsulated in the photothermal nanoparticles (Figure 1d). After immunostaining with a secondary Alexa Fluor-488-labeled anti-IgG2b antibody, subsequent visualization by fluorescence microscopy indicated specific staining of the neutralizing antibody-conjugated photothermal nanoparticles, as demonstrated by good colocalization between the Alexa Fluor-488 signal and the DiR dye (Pearson’s correlation coefficient ≈ 0.9, calculated by ImageJ).

Since the photothermal nanoparticles exhibited excellent absorption in the near-infrared region, we hypothesized that they could effectively generate local heating after light excitation with a suitable wavelength. To test this hypothesis, we applied a 650-nm LED with a power density of 250 mW/cm^2^ to excite the photothermal nanoparticles, because it would fit the absorption spectrum of the nanoparticles and could be safely used to excite healthy cells and tissues.^48, 49^ The nanoparticles were dispersed in phosphate buffered saline (1 × PBS; pH 7.4) at a concentration of 100 μg/mL, then subjected to LED excitation. As expected, a time-dependent temperature increase from 20 °C to 50 °C was observed within 10 min after light exposure (Figure 2a). In contrast, less than 2 °C of temperature change was observed in PBS alone, indicating the effect was specific to the nanoparticles (Figure 2a). Since the temperature of the entire solution exceeded 50 °C, we rationalized that the local heating rate of the photothermal nanoparticles should be much higher, and would thus enable effective inactivation of SARS-CoV-2. To test this hypothesis, we further evaluated local temperature changes following excitation using an LED-coupled voltage-clamp setup (Figure 2b) to understand the transient photothermal response of the nanoparticles.^50^ The nanoparticles were drop-casted onto the glass coverslip and the aggregate was covered with a PBS solution. The glass micropipette electrode was placed in close proximity to the surface of the aggregated nanoparticles. Then, a 650-nm LED was used to deliver 10 ms light pulses and the current response through the pipette was measured (Figure S2). The analysis revealed a 0.7 °C increase in the local temperature after each light pulse (Figure 2c), which corresponds to an initial heating rate of 70 °C/s. After the end of the pulse, the temperature decayed with a ~28 ms time constant. While the local temperature change observed in the measurement is lower than observed previously for silicon nanostructures,^51, 52^ the light density used for excitation is more than 100-times lower due to application of LED rather than focused laser. Additionally, the longer decay time constant suggests higher thermal resistance at the organic nanoparticle-water interface compared to inorganic nanostructures.^50^ The measurements further suggest strong photothermal response produced by the nanoparticles, which we expected would inactivate the viral particles due to the intense local heating.

**Figure 2.**
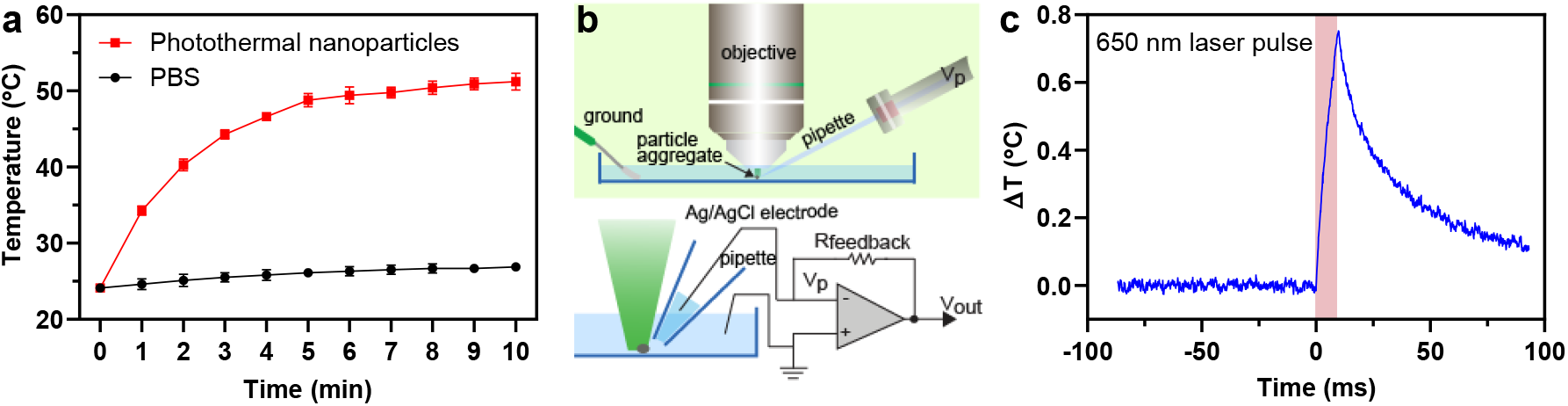
Photothermal characterization of the nanoparticles. **(a)** Temperature changes of the photothermal nanoparticles (200 μL) and 1 × PBS (200 μL) after the excitation with a 650-nm LED at the indicated times. Error bars indicate ± SEM. n = 3 per group. **(b)** Schematic diagram (top) and electrical diagram (bottom) of voltage clamp microscope setup used to measure local temperature change. **(c)** A representative trace of temperature increase during 10 ms excitation with a 650-nm LED at 1.7 W/cm^2^ intensity.

Next, we sought to determine if we could use the newly engineered photothermal nanoparticles to inhibit the infection of ACE2-expressing host cells by SARS-CoV-2. Due to the high risk of infection associated with live SARS-CoV-2, we decided to utilize alternative replication-incompetent viruses pseudotyped with the SARS-CoV-2 spike protein in order to evaluate efficacy of the nanoparticles. Pseudotyped viruses have been routinely employed to provide significantly safer conditions in which to study highly infectious viruses.^53–56^ As an initial proof-of-concept, we employed the vesicular stomatitis virus pseudotyped with the SARS-CoV-2 spike protein (SARS-CoV-2 VSV-GFP). The SARS-CoV-2 VSV-GFP pseudovirus enables transient expression of green fluorescent protein (GFP) upon entry into the host cell (Figure 3a), facilitating direct monitoring of viral uptake following incubation with the nanoparticles and photothermal treatment. In the absence of nanoparticles, HEK293T cells engineered to overexpress ACE2 (ACE2/HEK293T) were susceptible to pseudovirus infection as demonstrated by GFP expression after 24 h incubation (Figure 3b). A dose-dependent containment of the pseudovirus was observed upon addition of the antibody-conjugated photothermal nanoparticles (0.5 ~ 10 μg/mL) and 650-nm LED excitation (250 mW/cm^2^, 10 min). Complete inhibition of pseudovirus infection was achieved at a concentration of 5 μg/mL with an IC50 value of 1.68 μg/mL (Figure 3c, 3d and S3a), demonstrating the good virus inactivation efficiency of the nanoparticles. To confirm this effect was not dependent on the structure of the pseudovirus, we utilized a replication-incompetent lentivirus pseudotyped with SARS-CoV-2 spike protein (SARS-CoV-2 lentivirus-GFP), which stably integrates a gene encoding a GFP reporter into the host genome after infection (Figure 4a), for the subsequent in vitro viral infection study. As expected, in the absence of nanoparticles, the pseudovirus infected the ACE2/HEK293T cells, as indicated by GFP expression monitored at 48 h post infection (Figure 4b and S4a). After incubating with the neutralizing antibody-conjugated photothermal nanoparticles and applying 650-nm LED excitation (250 mW/cm^2^, 10 min), infection by the SARS-CoV-2 lentivirus-GFP was inhibited in a dose-dependent fashion when the concentration of nanoparticles was increased from 0.5 μg/mL to 10 μg/mL, demonstrating complete inhibition of pseudovirus infection at a concentration of 5 μg/mL and an IC50 of 0.23 μg/mL (Figure 4c, 4d, S3b, and S4b). Taken together, these data demonstrate the excellent inhibitory role of our antibody-photothermal nanoparticles in inactivating the SARS-CoV-2 pseudoviruses through the synergistic effect of the neutralizing antibody and photothermal treatment. Importantly, no toxicity was observed after treating the ACE2/HEK293T cells with different concentrations of the photothermal nanoparticles (Figure S5), providing further support for their safe use in biological systems.

**Figure 3.**
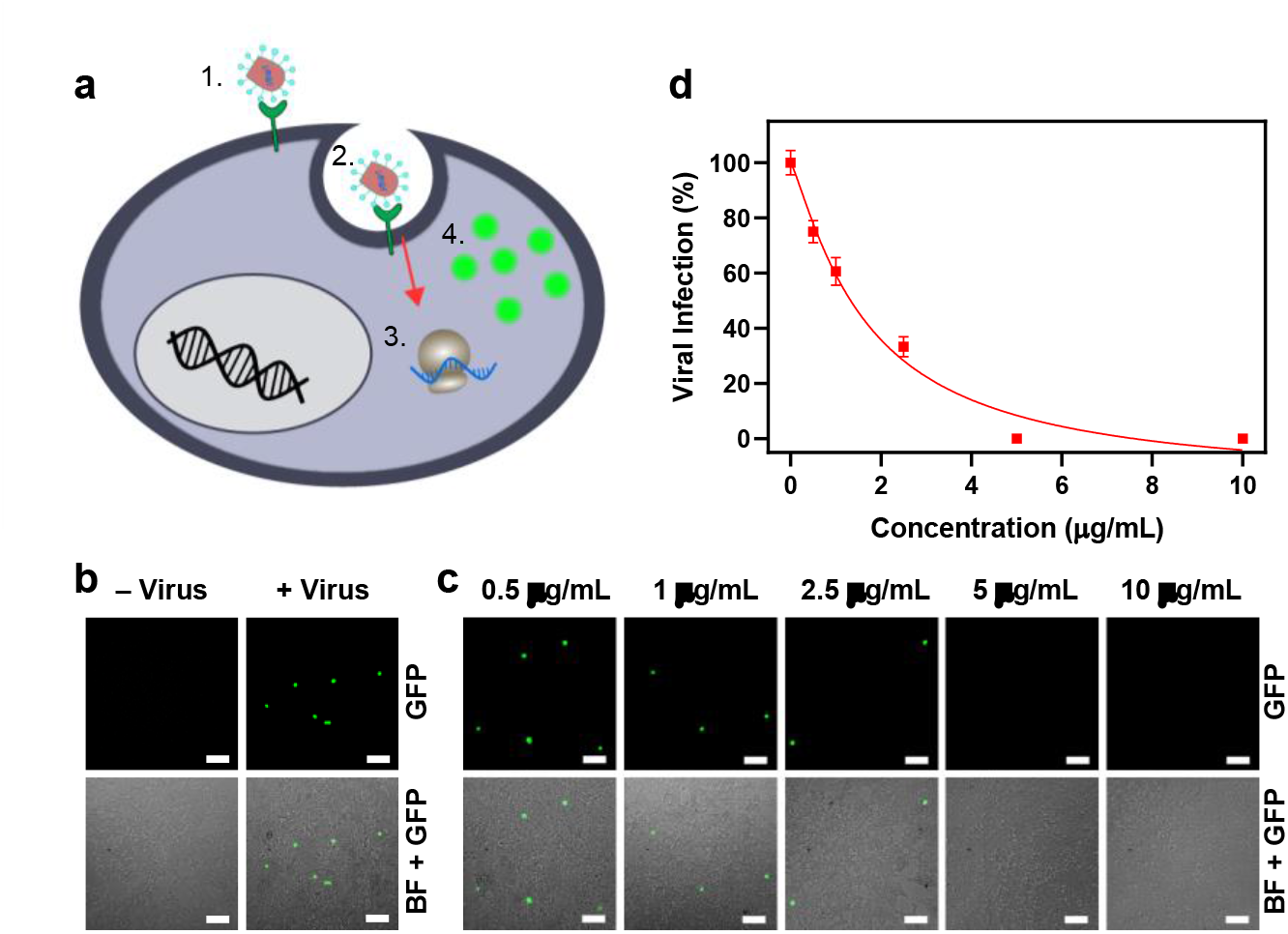
SARS-CoV-2 VSV-GFP pseudovirus infection in ACE2/HEK293T cells. (**a**) Schematic illustration of SARS-CoV-2 VSV-GFP pseudovirus infecting ACE2/HEK293T cells: the spike protein on the pseudotyped virus surface binds to ACE2 (1) and infects the cell (2), releasing its RNA to be transiently translated by the host (3) into GFP (4). **(b)** Fluorescent and brightfield (BF)-GFP merge images of ACE2/HEK293T cells before and after incubation with SARS-CoV-2 VSV-GFP. Scale bars: 100 μm. **(c)** Fluorescent and BF-GFP merge images of ACE2/HEK293T cells after incubation with SARS-CoV-2 VSV-GFP treated by different concentrations of the neutralizing antibody-conjugated photothermal nanoparticles and 650-nm LED excitation (250 mW/cm^2^, 10 min). Scale bars: 100 μm. **(d)** SARS-CoV-2 VSV-GFP infectivity after incubation with different concentrations of the neutralizing antibody-conjugated photothermal nanoparticles and 650-nm LED excitation (250 mW/cm^2^, 10 min). An IC50 of 1.68 μg/mL was calculated from dose-response curve. Error bars indicate ± SEM. n = 3 per group.

**Figure 4.**
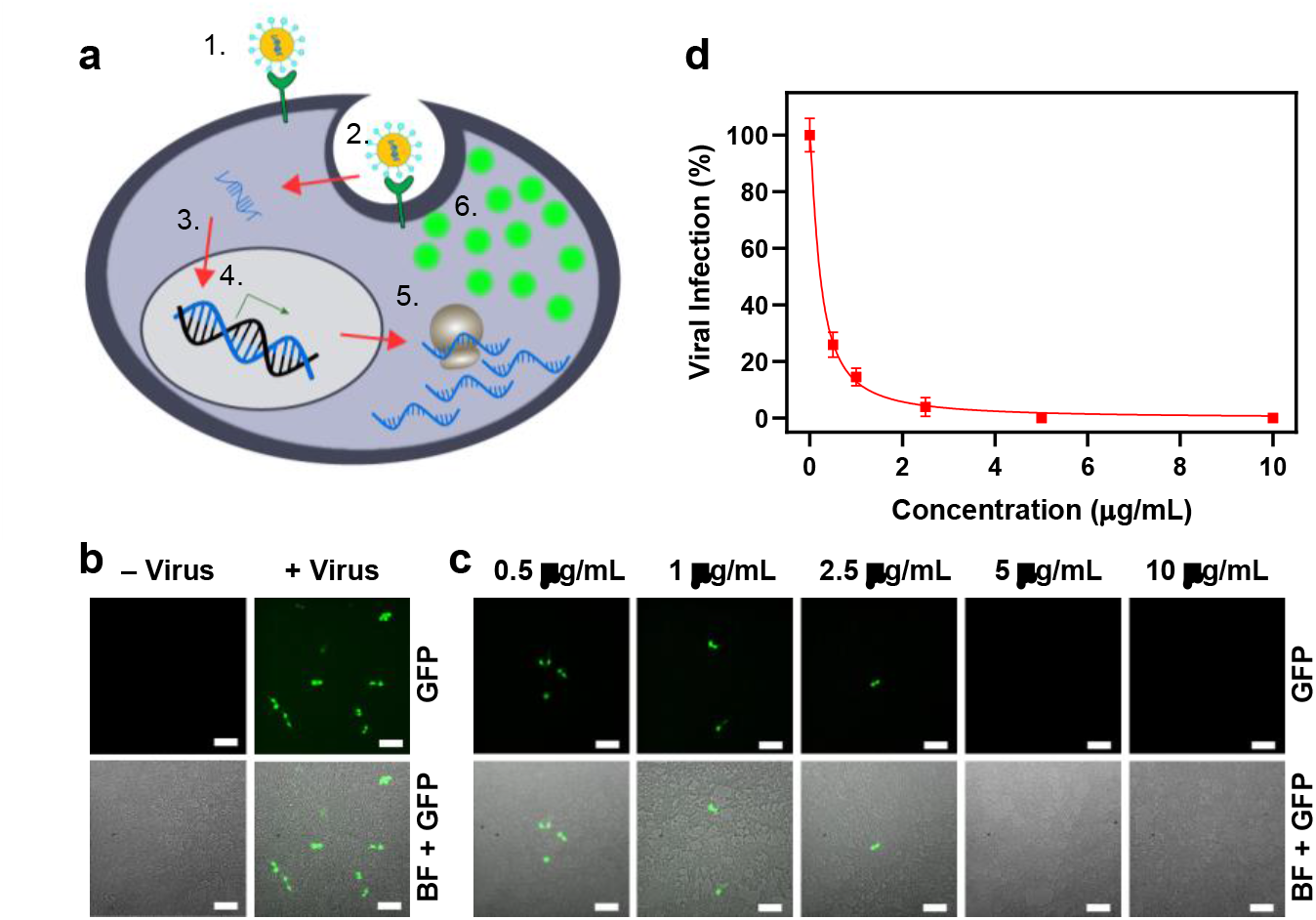
SARS-CoV-2 lentivirus-GFP pseudovirus infection in ACE2/HEK293T cells. (**a**) Schematic illustration of SARS-CoV-2 lentivirus-GFP pseudovirus infecting ACE2/HEK293T cells. The spike protein on the pseudotyped virus surface binds to ACE2 (1), and infects the cell (2). The viral RNA is integrated into the host DNA (3) where it constitutively transcribes (4) and translates it (5) into GFP (6). **(b)** Fluorescent and BF-GFP merge images of ACE2/HEK293T cells before and after incubation with SARS-CoV-2 lentivirus-GFP. Scale bars: 100 μm. **(c)** Fluorescent and BF-GFP merge images of ACE2/HEK293T cells after incubation with SARS-CoV-2 lentivirus-GFP treated by different concentrations of the neutralizing antibody-conjugated photothermal nanoparticles and 650-nm LED excitation (250 mW/cm^2^, 10 min). Scale bars: 100 μm. **(d)** SARS-CoV-2 lentivirus-GFP infectivity after incubation with different concentrations of the neutralizing antibody-conjugated photothermal nanoparticles and 650-nm LED excitation (250 mW/cm^2^, 10 min). An IC50 of 0.23 μg/mL was calculated from dose-response curve. Error bars indicate ± SEM. n = 6 per group.

## CONCLUSION

In summary, we developed an antibody-conjugated photothermal nanoparticle for active capture and inactivation of SARS-CoV-2. The nanoparticles displayed excellent photothermal properties as demonstrated by generation of a significant amount of local heat upon LED light excitation in a short time. In the virus neutralization experiments, the antibody-conjugated photothermal nanoparticles together with LED excitation enabled successful capture and inactivation of two different types of SARS-CoV-2 pseudoviruses, which resulted in complete protection against viral infection in ACE2/HEK293T cells. The synergistic effect of antibody capture and photothermal treatment provides an excellent virus neutralization efficiency. This work demonstrates a proof-of-concept that photothermal inactivation could potentially serve as a preventative approach to inhibit ADE that arises due to incomplete neutralization of the SARS-CoV-2 during SARS-CoV-2 infection and vaccination. Finally, our photothermal nanoparticle provides a flexible platform that can be readily adapted to target other strains of SARS-CoV-2 via conjugation to different antibodies or other novel therapeutic proteins and can be extended to other viruses, bacteria^57^ and malignant cells.^58^

## MATERIALS AND METHODS

### Preparation of Photothermal Nanoparticles

PCPDTBT (0.2 mg) and DSPE-PEG2000-NHS (2 mg) were dissolved and mixed in tetrahydrofuran (500 μL).^47^ For DiR labeled nanoparticles, 0.04 mg of DiR was further added into the mixture. The mixture was injected into 1 × PBS (10 mL) and sonicated using a probe sonicator (Qsonica Ultrasonic Homogenizer) on ice for 2 min. THF was then evaporated by rotary evaporator. The obtained nanoparticles were filtered through a 0.22 μm syringe filter and concentrated by ultrafiltration (Amicon^®^ Ultra-15 Centrifugal Filter Unit with 100 KDa cutoff, 4000 × g, 4 °C, 5 min). Anti-SARS-CoV-2 neutralizing antibodies were added into the nanoparticle solution with a mass ratio of 1:10 (antibody: DSPE-PEG2000-NHS) and stirred overnight at 4 °C to complete the conjugation.

### Nanoparticle Characterization

The measurement of UV-vis absorption spectra was carried out using a UV-vis spectrophotometer (Thermo Scientific NanoDrop™ 2000). The sizes of the nanoparticles were measured by dynamic light scattering (DLS) particle size analyzer (Malvern Zetasizer). The sizes and morphologies of the nanoparticles were studied by transmission electron microscopy (TEM, JEM-2010F, JEOL, Japan).

### Fluorescent Imaging of Photothermal Nanoparticles

The photothermal nanoparticles (100 μg/mL, 10 μL) was incubated with the secondary Alexa Fluor 488-labeled anti-IgG2b antibody (100 μg/mL, 10 μL) for 1 h. Single particle fluorescent images were captured by a fluorescence microscopy with a 100×/1.49 numerical aperture (NA) objective using a Nikon Ti-E inverted microscope. A 7-color solid state LED light source was connected with a liquid light guide to the microscope; the light then passes through a quad bandpass filter (ZET405-488-532-647m). The LED excitation wavelengths used were 470 ± 25 nm for green channel and 740 ± 20 nm for red channel (Spectra X, Lumencor). The emissions from the nanoparticles and Alexa Fluor-488 were captured by an Andor iXon Ultra 888 back-illuminated electron multiplying CCD (EMCCD) camera (Oxford Instruments).

### Photothermal Test of Photothermal Nanoparticles

Photothermal nanoparticles (100 μg/mL in 1 × PBS) were added into a well in a 96-well plate (flat bottom, GenClone^®^). A 650-nm LED (Spectra X, Lumencor) was applied to excite the nanoparticle solution for 10 min with a power density of 250 mW/cm^2^. The temperature changes from 0 to 10 min were recorded by a probe thermometer. Each measurement was repeated three times. Temperature changes of pure 1 × PBS were also measured using the same method.

### Measurement of Transient Photothermal Response of Photothermal Nanoparticles

Photothermal response measurements were performed on an upright microscope (Olympus, BX61WI) with a 20×/0.5 NA water-immersion objective. An LED light source (Lumencore Spectra X) with a 650 nm filter was used for excitation and was electronically controlled using transistor-transistor logic signals delivered from a digitizer (Molecular Devices, Digidata 1550). Voltage-clamp measurements were performed using Axopatch 200B amplifier (Molecular Devices). For the local temperature measurements, a pipette electrode (~0.8 MΩ at room temperature) filled with buffer solution (1 × PBS) was placed in a close proximity to the nanoparticles aggregated on top of glass coverslip submerged in PBS solution. Heat-induced currents were recorded in voltage-clamp mode. A laser pulse of 10 ms was delivered to the preparation 100 ms after the voltage jumped 0.5 mV below the holding potential. Current traces were recorded for the set of holding currents (Figure S2a) and the linear relationship between the holding current and photothermal current was established for different time points (Figure S2b). Temperature was inferred using a temperature-resistance calibration curve measured for the pipette.

### In Vitro Viral Infection of SARS-CoV-2 VSV-GFP Pseudovirus

ACE2/HEK293T cells (5 × 10^4^ cells) were seeded into 96-well plates (flat bottom, GenClone^®^) and incubated overnight at 37 °C in a humified CO2 incubator. A dilution series of the photothermal nanoparticles was prepared in a final volume of 50 μL of Dulbecco’s Modified Eagle Medium (DMEM) for a single well. Simultaneously, 15 μL of SARS-CoV-2 VSV-GFP (2 × 10^5^ titering units (TU)/mL) was diluted into DMEM to a final volume of 50 μL for a single well. The photothermal nanoparticles and SARS-CoV-2 VSV-GFP were combined to a total volume of 100 μL and incubated at 37°C for 1 h. After the initial incubation, the mixtures were excited by the 650-nm LED for 10 min with a power density of 250 mW/cm^2^ and added to cells. After 2 h incubation, 100 μL of fresh cell culture media were further added into each well. Each group contains three wells for the subsequent imaging and quantification. After 24 h incubation, the cells were imaged with Leica fluorescence microscope using 10×/0.30 NA objective. The LED excitation wavelengths were 470 ± 25 nm for GFP (Spectra X, Lumencor). The emissions were captured by an Andor iXon Ultra 888 back-illuminated EMCCD camera (Oxford Instruments). GFP-positive cells were counted manually three times by objective and the viral infection rates were calculated as the ratio of GFP-positive cells in the group incubated with the nanoparticles and virus to that of the group incubated with virus alone.

### In Vitro Viral Infection of SARS-CoV-2 Lentivirus-GFP Pseudovirus

ACE2/HEK293T cells (2 × 10^4^ cells) were seeded into 96-well plates (flat bottom, GenClone^®^) and incubated overnight at 37 °C in a humified CO2 incubator. A serial dilution of the photothermal nanoparticles was prepared in a final volume of 50 μL of cell culture media for a single well. Simultaneously, 10 μL of SARS-CoV-2 lentivirus-GFP (1 × 10^6^ TU/mL) was diluted into cell culture media to a final volume of 50 μL for a single well. The photothermal nanoparticles and SARS-CoV-2 lentivirus-GFP were combined to a total volume of 100 μL and incubated at 37°C for 1 h. After the initial incubation, the mixtures were excited by the 650-nm LED for 10 min with a power density of 250 mW/cm^2^ and added to cells. After 48 h incubation, cells were washed with 1 × PBS and fixed in 4% [w/v] paraformaldehyde for 15 min. Cells were then washed with 1 × PBS and further stained with Hoechst 33342 nuclear stain (2 μg/mL) for 15 min. The GFP expression monitored with a Nikon Ti-E inverted microscope using a 10×/0.30 NA objective. The LED excitation wavelengths were 395 ± 25 nm for Hoechst 33342 and 470 ± 25 nm for GFP (Spectra X, Lumencor). The emissions were captured by an Andor iXon Ultra 888 back-illuminated EMCCD camera (Oxford Instruments). The numbers of total cells and infected cells were counted in ImageJ using the following workflow: Adjust→Threshold (Otsu), Process→Binary→Fill Holes→Watershed, Analyze→ Analyze Particles. GFP expression was first normalized to total number of cells as calculated by DAPI staining. Then, viral infection rates were calculated as the ratio of normalized GFP positive cells in the group incubated with the virus and nanoparticles to that of the group incubated with virus only.

## ACKNOWLEDGMENTS

We thank Dr. Jeffrey Hubbell and his lab members for providing equipment for the nanoparticle synthesis. This work was supported by NIH New Innovator award 1DP2AI144245 (J.H.), NSF Career award 1653782 (J. H.) and NIDDK RC2DK122394 (E. C.). N.A. is supported by NIH T32DK007074 and J.R. is supported by the NSF Graduate Research Fellowships Program DGE-1746045.

## AUTHOR INFORMATION

### Author Contributions

X. C. and J. H. conceived the ideas and designed the project. J. H. supervised the project. X. C. synthesized, characterized the nanoparticles and conducted the in vitro viral infection experiments. A. P. measured the transient photothermal response and analyzed the data. Y. L. and J. S. performed the electron microscopy experiment and analyzed the data. N. A. and J. R. contributed to the imaging data analysis. M. C. maintained and provided the ACE2/HEK293T cells and advised on the in vitro SARS-CoV-2 VSV-GFP infection experiments. P. P.-M. produced the SARS-CoV-2 VSV-GFP pseudovirus and advised on the in vitro SARS-CoV-2 VSV-GFP infection experiments. X. C. and J. H. drafted the manuscript with input from A. P., N. A., J. R., E. C., and B. T.. E. C. and J. H. contributed to the funding acquisition.

### Notes

The authors declare no competing interests.

## REFERENCES

1. Ranney, M. L.; Griffeth, V.; Jha, A. K. Critical supply shortages—the need for ventilators and personal protective equipment during the Covid-19 pandemic. N. Engl. J. Med. (2020), 382, e41.

2. Spinelli, A.; Pellino, G. COVID-19 pandemic: perspectives on an unfolding crisis. Br. J. Surg. (2020), 707, 785–787.

3. Center for Systems Science and Engineering, Johns Hopkins University, COVID-19 Dashboard. 2020. https://coronavirus.jhu.edu/

4. Menni, C., et al. Real-time tracking of self-reported symptoms to predict potential COVID-19. Nat. Med. (2020), 26, 1037–1040.

5. Mehta, P.; McAuley, D. F.; Brown, M.; Sanchez, E.; Tattersall, R. S.; Manson, J. J.; Hlh Across Speciality Collaboration, U. K. COVID-19: consider cytokine storm syndromes and immunosuppression. Lancet (2020), 395, 1033–1034.

6. Zheng, Y.-Y.; Ma, Y.-T.; Zhang, J.-Y.; Xie, X. COVID-19 and the cardiovascular system. Nat. Rev. Cardiol. (2020), 17, 259–260.

7. Yan, R.; Zhang, Y.; Li, Y.; Xia, L.; Guo, Y.; Zhou, Q. Structural basis for the recognition of SARS-CoV-2 by full-length human ACE2. Science (2020), 367, 1444–1448.

8. Zhang, H.; Penninger, J. M.; Li, Y.; Zhong, N.; Slutsky, A. S. Angiotensin-converting enzyme 2 (ACE2) as a SARS-CoV-2 receptor: molecular mechanisms and potential therapeutic target. Intensive Care Med. (2020), 46, 586–590.

9. Jackson, L. A., et al. An mRNA Vaccine against SARS-CoV-2 — preliminary report. N. Engl. J. Med. 2020. DOI: 10.1056/NEJMoa2022483

10. Folegatti, P. M., et al. Safety and immunogenicity of the ChAdOx1 nCoV-19 vaccine against SARS-CoV-2: a preliminary report of a phase 1/2, single-blind, randomised controlled trial. Lancet (2020), 396, 467–478.

11. Le, T. T.; Andreadakis, Z.; Kumar, A.; Roman, R. G.; Tollefsen, S.; Saville, M.; Mayhew, S. The COVID-19 vaccine development landscape. Nat. Rev. Drug Discov. (2020), 19, 305–306.

12. Mulligan, M. J., et al. Phase I/II study of COVID-19 RNA vaccine BNT162b1 in adults. Nature 2020. https://doi.org/10.1038/s41586-020-2639-4

13. Mullard, A. COVID-19 vaccine development pipeline gears up. Lancet (2020), 395, 1751–1752.

14. Lurie, N.; Saville, M.; Hatchett, R.; Halton, J. Developing Covid-19 Vaccines at Pandemic Speed. N. Engl. J. Med. (2020), 382, 1969–1973.

15. Saini, K. S.; Lanza, C.; Romano, M.; de Azambuja, E.; Cortes, J.; de las Heras, B.; de Castro, J.; Lamba Saini, M.; Loibl, S.; Curigliano, G.; Twelves, C.; Leone, M.; Patnaik, M. M. Repurposing anticancer drugs for COVID-19-induced inflammation, immune dysfunction, and coagulopathy. Br. J. Cancer (2020), 123, 694–697.

16. Cohen, M. S. Hydroxychloroquine for the prevention of Covid-19 — searching for evidence. N. Engl. J. Med. (2020), 383, 585–586.

17. Wang, Y., et al. Remdesivir in adults with severe COVID-19: a randomised, double-blind, placebo-controlled, multicentre trial. Lancet (2020), 395, 1569–1578.

18. Shen, C., et al. Treatment of 5 critically Ill patients with COVID-19 with convalescent plasma. JAMA (2020), 323, 1582–1589.

19. Pérez-Cameo, C.; Marín-Lahoz, J. Serosurveys and convalescent plasma in COVID-19. EClinicalMedicine (2020), 23, 100370.

20. Arvin, A. M.; Fink, K.; Schmid, M. A.; Cathcart, A.; Spreafico, R.; Havenar-Daughton, C.; Lanzavecchia, A.; Corti, D.; Virgin, H. W. A perspective on potential antibody-dependent enhancement of SARS-CoV-2. Nature (2020), 584, 353–363.

21. Iwasaki, A.; Yang, Y. The potential danger of suboptimal antibody responses in COVID-19. Nat. Rev. Immunol. (2020), 20, 339–341.

22. Eroshenko, N.; Gill, T.; Keaveney, M. K.; Church, G. M.; Trevejo, J. M.; Rajaniemi, H. Implications of antibody-dependent enhancement of infection for SARS-CoV-2 countermeasures. Nat. Biotechnol. (2020), 38, 789–791.

23. Wan, Y.; Shang, J.; Sun, S.; Tai, W.; Chen, J.; Geng, Q.; He, L.; Chen, Y.; Wu, J.; Shi, Z.; Zhou, Y.; Du, L.; Li, F. Molecular mechanism for antibody-dependent enhancement of coronavirus entry. J. Virol. (2020), 94, e02015–19.

24. Brouwer, P. J. M., et al. Potent neutralizing antibodies from COVID-19 patients define multiple targets of vulnerability. Science (2020), 369, 643–650.

25. Cao, Y., et al. Potent Neutralizing antibodies against SARS-CoV-2 identified by high-throughput single-cell sequencing of convalescent patients’ B cells. Cell (2020), 782, 73–84.e16.

26. Rogers, T. F., et al. Isolation of potent SARS-CoV-2 neutralizing antibodies and protection from disease in a small animal model. Science (2020), 369, 956–963.

27. Wrapp, D.; Wang, N.; Corbett, K. S.; Goldsmith, J. A.; Hsieh, C.-L.; Abiona, O.; Graham, B. S.; McLellan, J. S. Cryo-EM structure of the 2019-nCoV spike in the prefusion conformation. Science (2020), 367, 1260–1263.

28. Lan, J.; Ge, J.; Yu, J.; Shan, S.; Zhou, H.; Fan, S.; Zhang, Q.; Shi, X.; Wang, Q.; Zhang, L.; Wang, X. Structure of the SARS-CoV-2 spike receptor-binding domain bound to the ACE2 receptor. Nature (2020), 587, 215–220.

29. Shang, J.; Ye, G.; Shi, K.; Wan, Y.; Luo, C.; Aihara, H.; Geng, Q.; Auerbach, A.; Li, F. Structural basis of receptor recognition by SARS-CoV-2. Nature (2020), 587, 221–224.

30. Wang, Q.; Zhang, Y.; Wu, L.; Niu, S.; Song, C.; Zhang, Z.; Lu, G.; Qiao, C.; Hu, Y.; Yuen, K.-Y. Structural and functional basis of SARS-CoV-2 entry by using human ACE2. Cell 2020.

31. Chen, X., et al. Human monoclonal antibodies block the binding of SARS-CoV-2 spike protein to angiotensin converting enzyme 2 receptor. Cell Mol. Immunol. (2020), 17, 647–649.

32. Wu, Y., et al. A noncompeting pair of human neutralizing antibodies block COVID-19 virus binding to its receptor ACE2. Science (2020), 368, 1274–1278.

33. Chi, X., et al. A neutralizing human antibody binds to the N-terminal domain of the Spike protein of SARS-CoV-2. Science (2020), 369, 650–655.

34. Liu, L., et al. Potent neutralizing antibodies against multiple epitopes on SARS-CoV-2 spike. Nature (2020), 584, 450–456.

35. Shi, R., et al. A human neutralizing antibody targets the receptor-binding site of SARS-CoV-2. Nature (2020), 584, 120–124.

36. Qiu, G.; Gai, Z.; Tao, Y.; Schmitt, J.; Kullak-Ublick, G. A.; Wang, J. Dual-Functional Plasmonic Photothermal Biosensors for Highly Accurate Severe Acute Respiratory Syndrome Coronavirus 2 Detection. ACS Nano (2020), 14, 5268–5277.

37. Shan, B., et al. Multiplexed Nanomaterial-Based Sensor Array for Detection of COVID-19 in Exhaled Breath. ACS Nano (2020), 14, 12125–12132.

38. Moitra, P.; Alafeef, M.; Dighe, K.; Frieman, M. B.; Pan, D. Selective Naked-Eye Detection of SARS-CoV-2 Mediated by N Gene Targeted Antisense Oligonucleotide Capped Plasmonic Nanoparticles. ACS Nano (2020), 14, 7617–7627.

39. Seo, G., et al. Rapid Detection of COVID-19 Causative Virus (SARS-CoV-2) in Human Nasopharyngeal Swab Specimens Using Field-Effect Transistor-Based Biosensor. ACS Nano (2020), 14, 5135–5142.

40. McKay, P. F., et al. Self-amplifying RNA SARS-CoV-2 lipid nanoparticle vaccine candidate induces high neutralizing antibody titers in mice. Nat. Commun. (2020), 11, 3523.

41. Zeng, C.; Hou, X.; Yan, J.; Zhang, C.; Li, W.; Zhao, W.; Du, S.; Dong, Y. Leveraging mRNA Sequences and Nanoparticles to Deliver SARS-CoV-2 Antigens In Vivo. Adv. Mater. (2020), 32, 2004452.

42. Medhi, R.; Srinoi, P.; Ngo, N.; Tran, H.-V.; Lee, T. R. Nanoparticle-based strategies to combat COVID-19. ACS Appl. Nano Mater. (2020), 3, 8557–8580.

43. Zhang, Q.; Honko, A.; Zhou, J.; Gong, H.; Downs, S. N.; Vasquez, J. H.; Fang, R. H.; Gao, W.; Griffiths, A.; Zhang, L. Cellular nanosponges inhibit SARS-CoV-2 infectivity. Nano Lett. (2020), 20, 5570–5574.

44. Weiss, C., et al. Toward Nanotechnology-Enabled Approaches against the COVID-19 Pandemic. ACS Nano (2020), 14, 6383–6406.

45. Huang, X.; Jain, P. K.; El-Sayed, I. H.; El-Sayed, M. A. Plasmonic photothermal therapy (PPTT) using gold nanoparticles. Lasers Med. Sci. (2008), 23, 217–28.

46. Jain, P. K.; Huang, X.; El-Sayed, I. H.; El-Sayed, M. A. Noble metals on the nanoscale: optical and photothermal properties and some applications in imaging, sensing, biology, and medicine. Acc. Chem. Res. (2008), 41, 1578–1586.

47. Cai, X.; Bandla, A.; Chuan, C. K.; Magarajah, G.; Liao, L.-D.; Teh, D. B. L.; Kennedy, B. K.; Thakor, N. V.; Liu, B. Identifying glioblastoma margins using dual-targeted organic nanoparticles for efficient in vivo fluorescence image-guided photothermal therapy. Mater. Horiz. (2019), 6, 311–317.

48. Avci, P.; Gupta, A.; Sadasivam, M.; Vecchio, D.; Pam, Z.; Pam, N.; Hamblin, M. R. Low-level laser (light) therapy (LLLT) in skin: stimulating, healing, restoring. Semin. Cutan. Med. Surg. (2013), 32, 41–52.

49. Sorbellini, E.; Rucco, M.; Rinaldi, F. Photodynamic and photobiological effects of light-emitting diode (LED) therapy in dermatological disease: an update. Lasers Med. Sci. (2018), 33, 1431–1439.

50. Jiang, Y.; Parameswaran, R.; Li, X.; Carvalho-de-Souza, J. L.; Gao, X.; Meng, L.; Bezanilla, F.; Shepherd, G. M.; Tian, B. Nongenetic optical neuromodulation with silicon-based materials. Nat. Protoc. (2019), 14, 1339.

51. Jiang, Y.; Li, X.; Liu, B.; Yi, J.; Fang, Y.; Shi, F.; Gao, X.; Sudzilovsky, E.; Parameswaran, R.; Koehler, K. Rational design of silicon structures for optically controlled multiscale biointerfaces. Nat. Biomed. Eng. (2018), 2, 508–521.

52. Jiang, Y., et al. Heterogeneous silicon mesostructures for lipid-supported bioelectric interfaces. Nat. Mater. (2016), 15, 1023–1030.

53. Nie, J., et al. Establishment and validation of a pseudovirus neutralization assay for SARS-CoV-2. Emerg. Microbes Infect. (2020), 9, 680–686.

54. Chen, Q.; Nie, J.; Huang, W.; Jiao, Y.; Li, L.; Zhang, T.; Zhao, J.; Wu, H.; Wang, Y. Development and optimization of a sensitive pseudovirus-based assay for HIV-1 neutralizing antibodies detection using A3R5 cells. Hum. Vaccines Immunother. (2018), 14, 199–208.

55. Logvinoff, C.; Major, M. E.; Oldach, D.; Heyward, S.; Talal, A.; Balfe, P.; Feinstone, S. M.; Alter, H.; Rice, C. M.; McKeating, J. A. Neutralizing antibody response during acute and chronic hepatitis C virus infection. Proc. Natl. Acad. Sci. U.S.A (2004), 101, 10149–54.

56. Wang, Y.; Cui, R.; Li, G.; Gao, Q.; Yuan, S.; Altmeyer, R.; Zou, G. Teicoplanin inhibits Ebola pseudovirus infection in cell culture. Antiviral Res. (2016), 125, 1–7.

57. Yougbaré, S.; Mutalik, C.; Krisnawati, D. I.; Kristanto, H.; Jazidie, A.; Nuh, M.; Cheng, T.-M.; Kuo, T.-R. Nanomaterials for the photothermal killing of bacteria. Nanomaterials (2020), 10, 1123.

58. Doughty, A. C. V.; Hoover, A. R.; Layton, E.; Murray, C. K.; Howard, E. W.; Chen, W. R. Nanomaterial applications in photothermal therapy for cancer. Materials 2019, 12, 779.

